# rxncon 2.0: a language for executable molecular systems biology

**DOI:** 10.1101/107136

**Authors:** Jesper C. Romers, Marcus Krantz

## Abstract

Large-scale knowledge bases and models become increasingly important to systematise and interpret empirical knowledge on cellular systems. In signalling networks, as opposed to metabolic networks, distinct modifications of and bonds between components combine into very large numbers of possible configurations, or microstates. These are essentially never measured in vivo, making explicit modelling strategies both impractical and problematic. Here, we present *rxncon* 2.0, the second generation rxncon language, as a tool to define signal transduction networks at the level of empirical data. By expressing both reactions and contingencies (contextual constraints on reactions) in terms of elemental states, both the combinatorial complexity and the discrepancy to empirical data can be minimised. It works as a higher-level language natural to biologists, which can be compiled into a range of graphical formats or executable models. Taken together, the rxncon language combines mechanistic precision with scalability in a composable and compilable language, that is designed for building executable knowledge bases on the molecular biology of signalling systems.

## 1 Introduction

The cellular regulatory networks monitor the state of a cell and its surroundings, and control key cellular processes such as metabolism, cell division and apoptosis. One of the main challenges of systems biology is to provide a mechanistic (as opposed to phenomenological) understanding of these networks in terms of their elementary building blocks: the reactions between and states of biological molecules.

Mechanistic understanding requires collection and integration of knowledge before actual model building and simulation. This in turn requires a language in which these tasks are natural from the biologist’s perspective and, additionally, one that allows the same knowledge base to be used in different modelling methods. To support comprehensive mechanistic models, this language need to be *expressive* - to precisely, at a given abstraction level, capture empirical knowledge; *scalable* - to allow large-scale models; *composable* - to support iterative and collaborative model development; and either *executable* or compilable into executable model code. These criteria can be used to evaluate any modelling language in terms of suitability for large-scale models and hence potential for genome-scale modelling.

The metabolic modelling community has set the gold standard for large-scale modelling. Based on a stoichiometric reaction definition, even genome-scale models can be built and analysed efficiently (Thiele and Palsson, 2010). Essentially, the network species are divided in disjunct mass pools (metabolites), and reactions are modelled as mass transfer between these pools. For mass transfer networks, such as metabolic networks, this is an appropriate abstraction and has hence been highly successful for model definition and simulation. However, the success of metabolic modelling relies on two fundamental features of mass transfer networks: First, the principal entity (mass) flowing through the network is conserved. Second, reactions turn over metabolites, making all reactions mutually exclusive at the level of individual metabolites. Neither of these features is true for signalling, and this has a strong impact on the suitability of these methods for modelling signal transduction – in particular at a large scale.

The regulatory networks process information. The information is encoded primarily in state changes of components, with no or only limited or local transfer of mass. Consequently, the assumptions underlying the metabolic modelling approaches are not valid or useful for the regulatory networks. First, the principal entity (information) flowing through these networks is not conserved, invalidating the basic assumption of constraint based analysis methods. Second, different reactions acting on the same signalling component are typically not mutually exclusive. Hence, signalling components can exist in multiple distinct states, and these states are important for information transfer. In experiments, we usually measure the state at a single residue (e.g. (un)phosphorylated) or domain (e.g. (un)bound to a specific ligand), and we refer to these non-disjunct states as elemental or macroscopic states (Conzelmann *et al*., 2008; Borisov *et al*., 2008; Tiger *et al.,* 2012; Creamer *et al.,* 2012). To simulate these systems with the mass transfer logic, we first need to create a system of disjunct microstates by specifying the state at each residue and domain for each model species. This is problematic for two reasons (Chylek *et al.,* 2015): First, we create a model with a different resolution than the underlying data, introducing ambiguity in data-model mapping. Second, we run into a combinatorial problem for all but the simplest signalling systems.

The problems posed by combinatorial complexity are well-known (Hlavacek et *al.,* 2003; Rother et *al.*, 2013) and overcoming them is one of the principal challenges in the field. *Scalability* is a fundamental problem in the description of cellular networks (Hlavacek and Faeder, 2009), in particular when aiming for genome-scale models (GSMs). The actual problem is twofold, first in the model formulation, and second in the model execution: even when the formulation of a model does not run into scalability issues, the execution or simulation might still be infeasible. While the problems with scalability and combinatorics are widely considered to be an intrinsic property of signalling per se, one can also understand it as trying to model signal transduction at the wrong abstraction level (Münzner et *al.*, 2017). Indeed, methods that do not inflate the complexity beyond empirical knowledge has been shown to scale efficiently to even large signal transduction networks (Faeder et *al.,* 2005, 2009; Danos and Laneve, 2004; Tiger et *al.*, 2012). These methods “trace out” the degrees of freedom that are unconstrained by empirical data.

The basic idea of these methods is to adapt the resolution of the network definition to that of empirical knowledge. They adhere to the “don’t care, don’t write” principle. In the rule based formalisms, reactions consume or produce partially defined reactants and products that match sets of microstates (Faeder *et al.* (2005, 2009); Danos and Laneve (2004), reviewed in Chylek *et al.* (2015)). Hence, a rule-based approach allows for efficient model definition. To simulate these models, either the rules are used to generate the full set of microstates and reactions between them, which only works for small models, or the rules are used for stochastic, network-free simulation (Danos and Laneve, 2004; Faeder *et al.,* 2009; Sneddon *et al.,* 2011). With the reaction-contingency (rxncon, “reaction-con”) language, we describe the mechanistic building blocks responsible for cellular signalling: molecular *components* with their *elemental states,* that describe modifications of a single molecule or complexations between molecules, and *elemental reactions,* that describe independent reaction events, and their *contingencies,* the necessary molecular context consisting of interactions and modifications. The elemental reactions create and destroy elemental states, which in their turn make up the contingencies. In this sense, the language is very close to *experiment,* since every statement corresponds to an experimental fact. This makes the language composable, as single reaction events can be added to a system by adding single statements and without touching any previous statements. Similarly, as knowledge about a signalling network progresses, more accurate domain or residue information can be provided without having to completely start from scratch. The local impact of changes and extensions facilitates *iterative* model building and *cooperative* efforts by multiple groups. While rule based models are directly executable, rxncon networks are compilable into executable model code – including rule based models – giving the flexibility of multiple output formats. However, the first version of the rxncon language had limitations in expressiveness (Tiger *et al.,* 2012), for example related to the structure of larger molecular complexes and the mutual exclusivity of elemental states.

Here, we present the second generation rxncon language. We present a formal syntax and semantics for this thoroughly reworked language, and show that we address previous limitations in expressiveness and compilability. In particular, we added *structure indices* to unambiguously describe complexes with multiple identical subunits, developed the notion of *skeleton rules* to give semantics to elemental reactions, introduce explicit *reverse reactions* for reactions that are inherently bidirectional – which allows one to add different contingencies depending on the direction of the reaction, added explicit *neutral states* and introduce the notion of *mutual exclusivity* of states, which is closely related to the concept of *elemental resolution.* We demonstrate the improved expressiveness by translating an extensive and well annotated model of the pheromone response in yeast, one of the most well understood eukaryotic signalling pathways. The *rxncon* language is agnostic to the actual modelling method used to simulate the system under study. In this sense, it can be (loosely) compared to a higher-level computer programming language that has different “compilation targets”. Currently the language can be “compiled” to a Boolean network (Thieme *et al.,* 2017) or a rule-based model (Romers *et al.,* in preparation), and other targets are being studied. This means the modeller can work at the appropriate abstraction level to her (elemental reactions and elemental states) and leave it to a machine to provide the actual error-prone translation into a modelling formalism.

Taken together, we present a scalable, composable language for describing cellular signal transduction processes that contains the appropriate abstractions to make a direct connection with experimental knowledge and which is compilable to executable, simulatable models.

## 2 Syntax and semantics of the *rxncon* language

A *rxncon* system can be thought of as a compendium of knowledge about the mechanistic processes that underlie cellular signal transduction phenomena. The language, which we formalize below, consists of a collection of statements enumerating the biochemical reactions and their contingencies, the context in which these reactions take place.

Since each statement considers either a *reaction* or a *contingency*, each individual statement is an experimentally verifiable fact about the signalling network, that can be annotated with literature sources and further details. These statements are independent: the reactions only denote which property of a molecule (phosphorylation residue, binding domain) changes, without having to resort to a microstate description, which is inherently unscalable.

In these sections, we give a formal definition of the *rxncon* language. We use Backus-Naur Form (BNF) (Backus, 1959; Naur, 1961) definitions to describe syntactically correct *rxncon* statements, and show how the

BNF products map to different semantic concepts. These concepts in turn map to classes in the code of our implementation of the language. We will sometimes refer to a property p of an object by writing 〈object〉.p: by this we mean that part of the BNF product with the name p. Terms in square brackets are optional and the Kleene star ‘*’ means zero or more times, whereas ‘+’ means one or more times.

### 2.1 Specs

The central building block of the *rxncon* language is a molecule specification or *spec,* of which the BNF definition is given in (1). They appear as elements of reaction and state statements, in which they are used to specify properties of molecules.

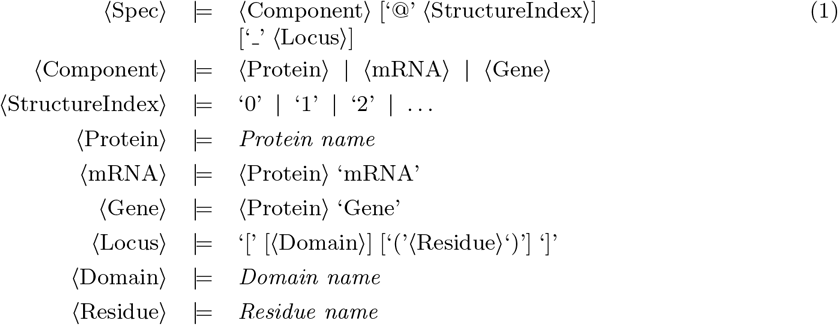

The required *Component* denotes the particular protein, gene or mRNA that is referred to. Protein names are composed of alphanumeric characters, but have to start with a letter and not end in –Gene or –mRNA, which automatically refer to the gene or mRNA molecule corresponding to the protein. This one-to-one-to-one relation makes implementation of reactions that rely on the central dogma, *i.e.* translations and transcriptions, straightforward.

The optional *Locus* points to a location on a molecule, in order of increased resolution: to a *domain* or a *residue*. Domains can contain residues. This construction allows one to accurately reflect the detail of experimental knowledge: *e.g.* one might not know the precise residue at which a protein needs to be phosphorylated in order for a certain reaction to be possible, but only the domain on which the phosphorylation lives. If a residue is specified, the spec’s *resolution* is “at the residue level”, and similar for domain. If no Locus is provided, the spec’s resolution is “at the component level”.

Larger molecular complexes that can appear in contingencies might have multiple subunits containing the same molecule. In such a case, there is an ambiguity when combining different contingencies based solely on the component names of the molecules. To work around this problem, we introduce an additional *Structure index*, a number unique for each molecule.

Specs have a superset / subset relation amongst each other. The spec A is a subset of a spec B if

- *A*’s and *B’s* Component and StructureIndex match, and
- *A*’s resolution is equal or higher than *B*’s, and
- the Locus information in *B* that is not empty coincides with that in A.

The spec *A* is a superset of a spec *B* if *B* is a subset of *A*. Trivially a spec is its own superset and its own subset.

### 2.2 States

States correspond to independent *observable quantities,* such as protein’s phosphorylation or bond to another protein. What is called “state” in the literature often refers to the fully specified microstate of a molecule. A *rxncon* state is a macroscopic state: except for the information on *e.g.* a phosphorylation, all other information is ignored (or “traced out” in statistical physics parlance).

In this section we discuss the different properties that states can have, and the different classes of states that appear in the *rxncon* language.

States belong to a certain *Class*. Currently we distinguish six classes in *rxncon*, see Table 1. *Modifications* such as `A_[(r)]-{p}` denote a modification of a particular residue, such as phosphorylations. *Interactions* such as `A_[a]--B_[b]` describe bound states between different molecules. *SelfInteractions* such as `A-[x]--[y]` describe bound states within the same molecule. *EmptyBindings* such as `A_[x]--0` describe an unbound (empty) binding domain on a molecule. *Inputs* such as `[Turgor]` describe a macroscopic input signal that cannot be localised on a single molecule. The special *FullyNeutral* state will be discussed below.

**Table 1:**
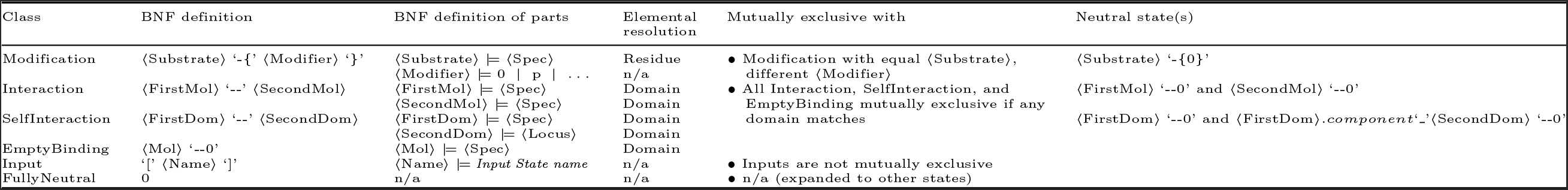
Classes of states in the *rxncon* language. We denote the BNF definition, the resolution of the respective specs and loci at which the state becomes elemental, the states with which they are mutually exclusive and their neutral counterparts.

We distinguish states that are *located* on a single molecule, Modifications, Selfinteractions and EmptyBindings, ones that are located on a pair of molecules, Interactions, and non-localizable Inputs.

States are built up of zero or more specs or loci and inherit the notion of resolution from them. Each class of state has for every spec or Locus an associated *elemental resolution*. If every spec and locus that appears in a state is at its elemental resolution, the state itself is referred to as an *elemental state*.

States inherit the superset / subset relation from the specs they contain. A state S_1_ is a subset of a state S_2_ if

- they belong to the same class, and
- all non-spec properties coincide, and
- all specs in S_1_ are subsets of the specs in S_2_.

For classes of states that contain more than one spec, we consider the meaning of two states to coincide under permutation of the specs, *i.e.* `A_[a]--B_[b]` is equivalent to `B_[b]--A_[a]`.

There exists a notion of *mutual exclusivity* of states: the same residue on the same molecule cannot simultaneously be in the phosphorylated and unmodified form. An overview of which states are mutually exclusive with which can be found in Table 1. Note that elementarity of states is assumed here.

Every state has one or more “neutral” counterparts, for Modifications this is a Modification with the neutral Modifier, and for (Self)Interactions the appropriate EmptyBindings. Reactions that synthesise components mostly do so in a fully neutral combination of states, the FullyNeutral-State which we denote by “0”. This state is in fact a shorthand for the combination of all the neutral states for a particular component, see Section 2.6.

### 2.3 Syntax of reactions

The states we have seen in the previous section are created and destroyed by *elemental Reactions.* The syntax, which is presented in (2), contains two specs and a ReactionType.

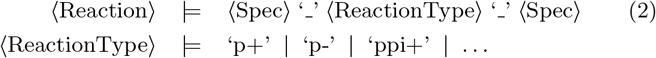

For a (non-exhaustive, but representative) list of *rxncon* Reactions, see Table 2. The skeleton rule that determines the semantics is explained in the following section.

**Table 2:**
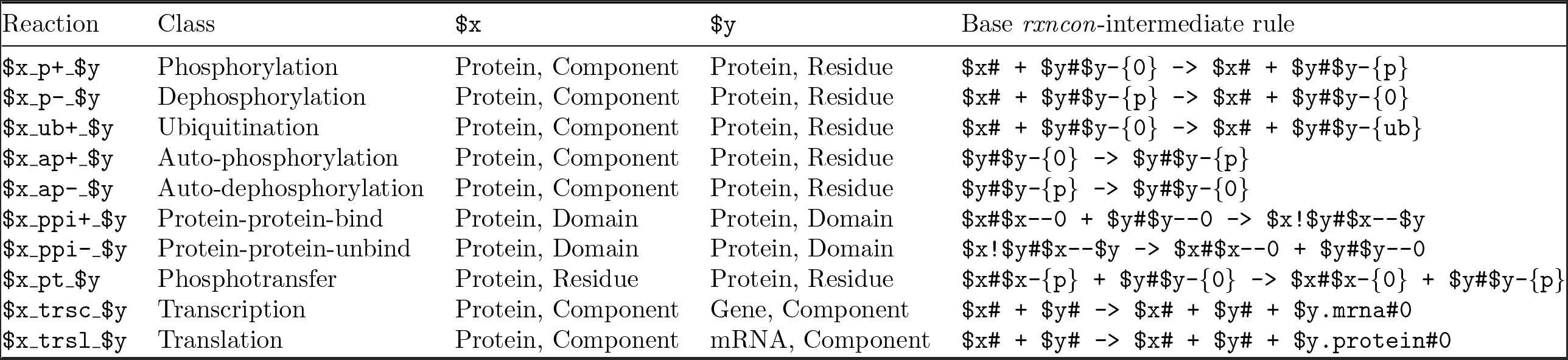
Classes of Reactions in the *rxncon* language. Here we denote the specs appearing in the Reaction as computer-code style variables **$x, $y** to allow a “method call” syntax on them which extracts a particular part of their BNF expression. For the domain-specific language expressing the rules, please see the text.

### 2.4 Semantics of reactions: skeleton rules

Several languages, such as the BioNetGen language (BNGL) Faeder *et al.* (2009) and Kappa (Danos *et al.,* 2007) exist to formulate rule-based models. Here we briefly define the skeleton rule language: a simple language that is used to define the semantics of the *rxncon* reactions in terms of previously introduced *rxncon* concepts.

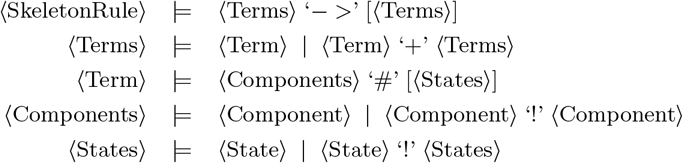

The rule describes a transition between one or more terms at its left-hand side into zero (in the case of decay for example) or more terms at its right-hand side. Every term consists of (i) one or more *Components,* which are connected in a complex and (ii) zero or more elemental states. The latter define the internal state of the molecules in the complex.

Given a Reaction and its skeleton rule, one can define the notions of *production*, *consumption*, *synthesis* and *degradation* of states by Reactions, where RHS and LHS refer to the right-hand side and left-hand side of the corresponding skeleton rule:

- a state is produced by a reaction if it appears on the RHS, not on the LHS, but the component carrying the state does appear on the LHS,
- a state is consumed by a reaction if it appears on the LHS, not on the RHS, but the component carrying the state does appear in the RHS,
- a state is synthesised by a reaction if it appears on the RHS, and the component carrying the state does not appear on the LHS,
- a state is degraded by a reaction if the component carrying the state appears on the LHS, no state mutually exclusive with it appears on the LHS, and the component carrying the state does not appear on the RHS.

### 2.5 Contingencies

The context for reaction events is given by *contingencies,* see (3). These are (Boolean combinations of) states that influence the reaction events. We refer to sections 2.2 and 2.4 for the production rules for states and skeleton rules (or reactions) respectively.

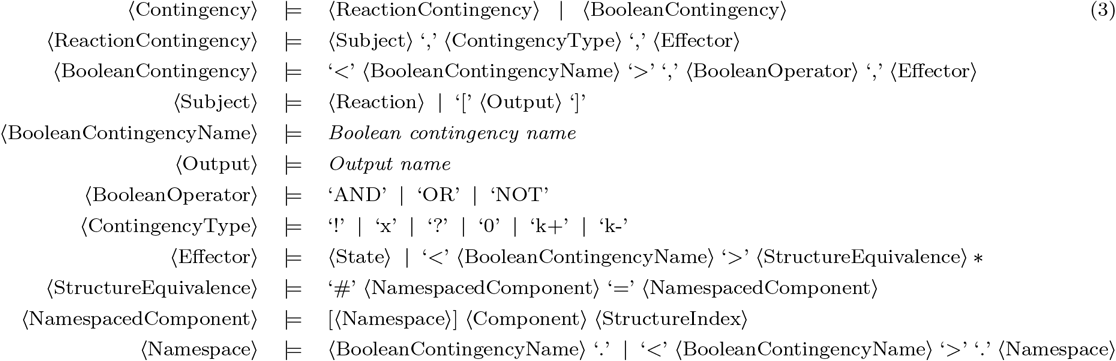

First, we distinguish between reaction contingencies and “Boolean” contingencies. The simplest example to understand is the reaction contingency consisting of the triple *reaction, contingency type* and *state,* in which a single state directly influences a reaction. Contingency types can be either *strict contingencies,* with “!” denoting an absolute requirement and “x” an absolute inhibition, or *quantitative contingencies,* with contingency type “k+” representing a positive contribution to the reaction rate and contingency type “k-” a negative contribution.

Finally, the contingency types “0” and “?” denote *no effect* respectively *unknown effect.*

Boolean contingencies can be used to describe more complex contexts, in which the combination of a number of states influences a reaction. The format described in (3) allows one to formulate arbitrarily nested Boolean expressions. In order for a boolean contingency to make sense, all contingencies carrying the same name must have the same Boolean operator.

When all of a reaction’s contingencies are satisfied, the signalling network is considered to be in a state that can accomodate the reaction. For a reaction to be considered active, the network needs to be in this state, the reaction’s reactants need to be present and its sources (the states it targets for consumption) need to be present.

Contingencies inherit the notion of elementarity from the states they contain: if all states are elemental, the contingency is elemental and otherwise not.

#### 2.5.1 Satisfiability of contingencies

Since contingencies can form Boolean expressions of states, it is important that they are satisfiable. The reference implementation of *rxncon* is linked to picoSAT (Biere, 2008), an industrial-strength satisfiability solver.

Every contingency can (and will, in practice) be expanded into an elemental contingency (see Section 2.6). It is therefore sufficient to consider satisfiability of elemental contingencies. However, not every naively obtained solution to a Boolean expression over states is a valid solution: some states are mutually exclusive with one another, and are therefore not allowed.

Furthermore, a contingency needs to be *connected* to the reactants: if it refers to a molecule that is not one of the reactants there needs to be at least one path from the reactants to that molecule over bond states to be valid. This in particular becomes an issue when translating a *rxncon* system to rules in a rule-based model (Romers *et al.,* in preparation). A Boolean contingency is *satisfiable* if it has at least one solution that contains no mutually exclusive states and is connected.

#### 2.5.2 Structured indices and boolean contingencies

In many cases, the name of a molecule might not be sufficient to uniquely identify it in a complex, which is solved by adding structure indices to specs. The *rxncon* reference implementation has an algorithm to find reasonable default structure indices if none are supplied, and internally every spec in the contingency list carries a structure index once the *rxncon* system has been constructed.

When one defines contingencies that contain Boolean expressions or nested Booleans (Boolean contingencies containing Boolean contingencies), there is an additional ambiguity. The structure indices of a Boolean contingency live in a namespace that is labelled by the name of that particular Boolean contingency. Within that namespace every structure index is well-defined, but one has to map the indices within the namespace of the Boolean contingency to the subject namespace. This applies to contingencies that have a reaction as their subject as well as contingencies that themselves have a Boolean contingency as their subject: when one combines multiple Boolean contingencies, the namespaces have to be merged to obtain an unambiguous labelling.

The following rules apply:

- for monomolecular reactions, the reactant has structure index 0,
- for bimolecular reactions, the reactants have indices 0 and 1,
- when a contingency (with a reaction or a Boolean contingency as its subject) has a Boolean contingency as its object, a *structure equivalence* has to be supplied. This equivalence relation establishes which (Component, StructureIndex) pairs in the subject namespace map to which (Component, StructureIndex) pairs in the object namespace. As an example, the equivalence #A@0=A@2 means that the component A@0 in the subject’s namespace refers to the same molecule as A@2 in the Boolean contingency’s namespace.

### 2.6 *rxncon* system

A full *rxncon* system is a set of one or more Reactions and zero or more Contingencies, see (4).

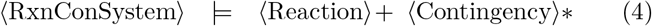

After reading a *rxncon* system, one first *finalizes* the system an then validates it.

The finalization concerns (1) the expansion of non-elemental contingencies and (2) the structuring of non-structured contingencies. The first happens in two places. It is possible to formulate contingencies in terms of non-elemental states, whereas Reactions by definition only produce, consume, synthesise and degrade elemental states. To handle this mismatch, every non-elemental state appearing in a contingency becomes a Boolean ‘OR’ complex carrying the name of the non-elemental contingency.

Furthermore, the FullyNeutral state needs to be expanded into a fully specified microstate. This state appears in synthesis reactions and is useful since it is a property of the entire *rxncon* system what the neutral state for a component exactly is.

Finally, a validation takes place. Not every *rxncon* system is internally consistent. However, this can only be decided after finalisation. The validation checks that

- there are no elemental states appearing in the contingencies that are not produced, consumed, synthesised or degraded by elemental reactions,
- there are no reactions that are the subject of contingencies that are not in the list of reactions,
- there are no unsatisfiable contingencies.

## 3 Language for executable biology: translating the HOG pathway

The *rxncon* language has been designed to be close to empirical molecular biology, which facilitates formalising empirical knowledge. Here we will illustrate this process by building a small model of a signal transduction pathway. As example we chose the well characterised High Osmolarity Glycerol (HOG) MAP kinase pathway of baker’s yeast, which is described clearly and concisely in (Hohmann, 2009). For this illustration, we only consider one of two input branches, the Slnl-Ypdl-Ssk1 phosphorelay system, and the Ssk2-Pbs2-Hog1 MAP kinase cascade. We consider activated Hog1 to be the output of the system. Reading through the first section on “The yeast HOG pathway”, we find the following sentences, which we translate to *rxncon* statements:

- “Active Slnl is a dimer that performs auto-phosphorylation on a histidine.” There exists a dimerization reaction, in *rxncon* parlance, a protein-protein interaction (ppi) between two Sln1 proteins, the product state of which is a requirement (–) for its autophosphorylation (AP+) reaction:

~~~
- Reaction: Sln1_[Sln1]_ppi_Sln1_[Sln1]
~~~

~~~
- Reaction: Sln1_AP+_Sln1_[(His)]
~~~

~~~
- Contingency: Sln1_AP+_Sln1_[(His)],!,Sln1@0_[Sln1]–Sln1@2_[Sln1]
~~~

- “Slnl is active under ambient conditions and inactivated upon hyperosmotic shock.” The activity of Sln1 is furthermore regulated by an input state: the absence of hyperosmotic shock or, equivalently, the presence of Turgor pressure:

~~~
- Contingency: Sln1_AP+_Sln1_[(His)],!,[Turgor]
~~~

- “This phospho group is then transferred to a receiver domain in Sln1, further to Ypd1 and eventually to the receiver domain in Ssk1.” A cascade exists of three phospho-transfer (PT) reactions:

~~~
- Reaction: Sln1_[(His)] _PT_Sln1_[(Rec)]
~~~

~~~
- Reaction: Sln1_[(Rec)] _PT_Ypd1_[(P)]
~~~

~~~
- Reaction: Ypd1_[(P)] _PT_Ssk1_[(Rec)]
~~~

- “Phospho-Ssk1 is intrinsically unstable or dephosphorylated by an unknown phosphatase.” We introduce an auxiliary molecule Up1 (for Unknown Phosphatase), responsible for the dephosphorylation (P-) of the Sskl protein:

~~~
- Reaction: Upl_P-_Sskl_[(Rec)]
~~~

- “Ssk1 binds to the regulatory domain of the Ssk2 and Ssk22 MAPKKKs, which allows Ssk2 and Ssk22 to autophosphorylate and activate themselves.” The product state of the protein-protein interaction between Ssk1 and Ssk2 is a strict requirement for the autophosphorylation of Ssk2. In what follows we omit Ssk22, all statements are symmetric under exchange of Ssk2 with Ssk22.

~~~
- Reaction: Sskl_[Ssk2]_ppi_Ssk2_[Sskl]
~~~

~~~
- Reaction: Ssk2_AP+_Ssk2_[(auto)]
~~~

~~~
- Contingency: Ssk2_AP+_Ssk2_[(auto)],!,Sskl_[Ssk2]--Ssk2_[Sskl]
~~~

- “Phospho-Ssk1 is the inactive form and hence does not activate the downstream MAP kinase cascade.” The phosphorylated state of Ssk1 blocks the cascade. Since the only influence Ssk1 has on the pathway is exerted through its interaction with Ssk2, we choose to block (x) that particular reaction:

~~~
- Contingency: Sskl_[Ssk2] _ppi_Ssk2_[Sskl],x,Ssk1_[(Rec)]-{P}
~~~

- “Active Ssk2 and Ssk22 then phosphorylate and activate Pbs2, which in turn phosphorylates (on Thrl74 and Tyrl76) and activates Hogl.” The three reactions speak for themselves and the first three contingencies define the order in which they can take place. The last two contingencies define what we mean by the output of this pathway: doubly-phosphorylated Hogl on residues Thrl74 and Tyrl76.

~~~
- Reaction: Ssk2_P+_Pbs2_[(P)]
~~~

~~~
- Reaction: Pbs2_P+_Hogl_[(Thrl74)]
~~~

~~~
- Reaction: Pbs2_P+_Hogl_[(Tyrl76)]
~~~

~~~
- Contingency: Ssk2_P+_Pbs2_[(P)],!,Ssk2_[(auto)]-{P}
~~~

~~~
- Contingency: Pbs2_P+_Hogl_[(Thrl74)],!,Pbs2_[(P)]-{P}
~~~

~~~
- Contingency: Pbs2_P+_Hogl_[(Tyrl76)],!,Pbs2_[(P)]-{P}
~~~

~~~
- Contingency: [Output],!,Hogl_[(Thrl74)]-{P}
~~~

~~~
- Contingency: [Output],!,Hogl_[(Tyrl76)]-{P}
~~~

- We turn the page, skip the section on the Sho1-branch, but find finally find: “The phosphorylation state of the MAPK Hog1 is controlled by various protein phosphatases. Those include the phospho-tyrosine phosphatases Ptp2 and Ptp3 [2224] as well as the phospho-threonine phosphatase Ptc1”

~~~
- Reaction: Ptc1_P-_Hog1_[(Thr174)]
~~~

~~~
- Reaction: Ptp2_P-_Hog1_[(Tyr176)]
~~~

~~~
- Reaction: Ptp3_P-_Hog1_[(Tyr176)]
~~~

The full model is available from our model repository (https://github.com/rxncon/models), file HūG_example.xls, and is represented visually in Figure 1.

**Figure l:**
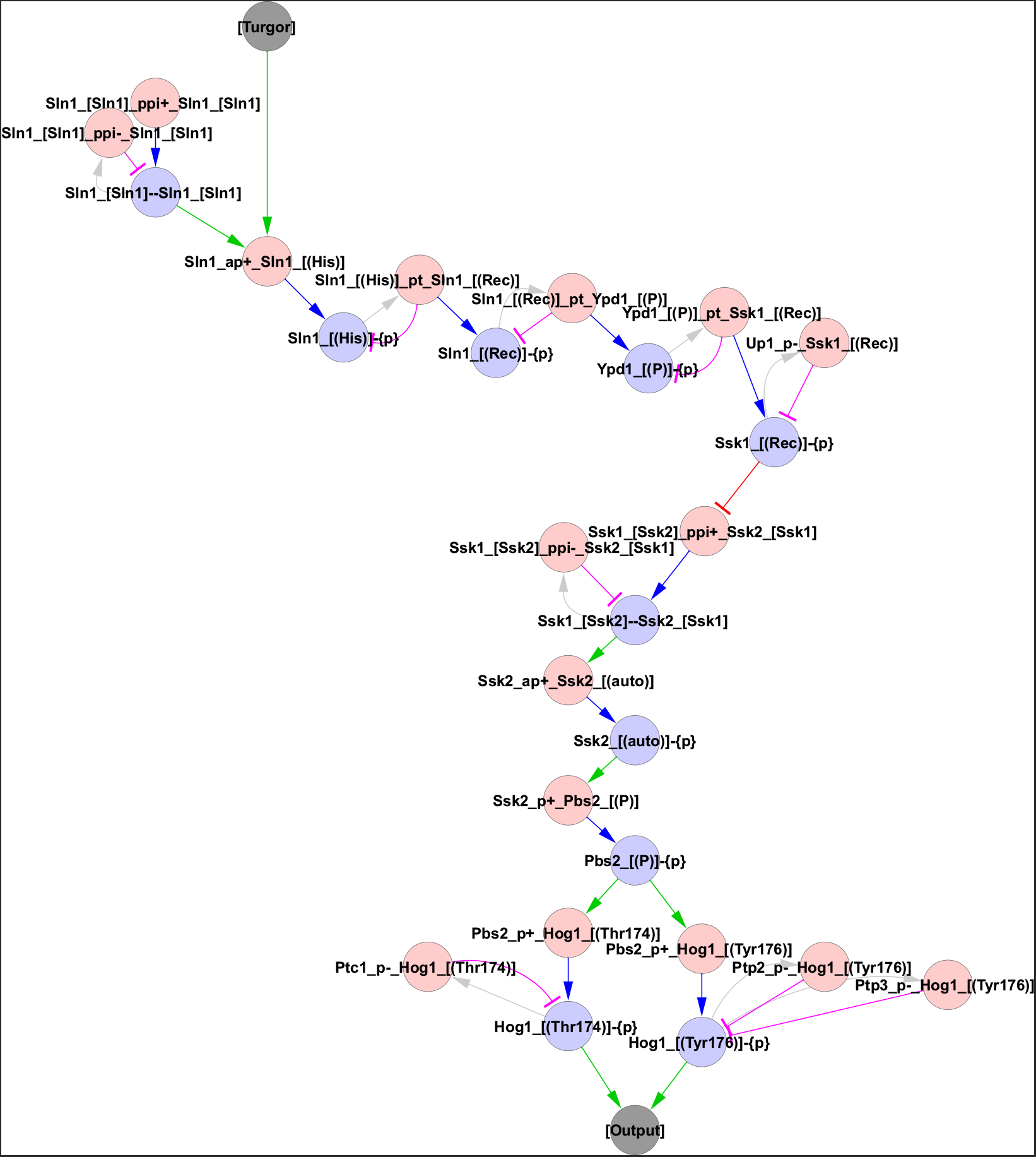
The HOG pathway model visualised in a regulatory graph. This bipartite graph displays the information flow over elemental reactions (red nodes) and elemental states (blue nodes). Elemental reactions produce (blue edges) or consume (purple edges) elemental states, which in turn act as source states (grey edges), inhibitors (red edges; x/k-contingencies) or activators (green edges; !/k+ contingencies) of elemental reactions. Input and output nodes (grey) indicate the model boundaries and act as states and reactions, respectively. The regulatory graph clearly visualises both the model assumptions (each elemental reaction and contingency is displayed) and the information flow through the network. Here, turgor activates Slnl homodimers to autophosphorylate on the His residue (as both the input and the dimerisation are required for the autophosphorylation), which provide the initial source state for the phos-photransfer chain through the receiver domain of Slnl on to Ypdl and finally Sskl. The Sskl phosphorylation in turn inhibits the downstream MAP kinase cascade by preventing the `Ssk1—Ssk2` dimerisation which is required for Ssk2 autophosphorylation. When this inhibition is relieved, the autophosphorylation triggers the kinase cascade: It allows Ssk2 to phosphorylate Pbs2, which in turn is required to phosphorylate Hogl on two sites its activation loop. Dually phosphorylated Hogl is active, triggering the pathway outputs. Hence, the information path can be followed from the input to the output via directed edges. All information is taken from Hohmann (2009), see text for details.

## 4 Scalability and expressiveness: translating the yeast pheromone pathway

To examine the scalablility and expressiveness of *rxncon* 2.0, we translated a rule based model of the yeast pheromone response pathway to the rxncon language. This is one of the largest and most well annotated rule based models that we are aware of, and it defines a microstate system of over 200.000 states which the authors consider too complex for (meaningful) simulations (Thomson *et al.,* 2011) (http://yeastpheromonemodel.org/wiki/Extracting_the_model). Hence, it is a suitable target to analyse scalability. In addition, it was a challenging target for the first version of rxncon, where we for example failed to express trans-phosphorylation across homodimer scaffolds (Tiger *et al.,* 2012). Taken together, the model describes a medium sized signalling pathway at mechanistic resolution with several challenging features (combinatorics, complexes, homodimers), providing a suitable benchmark for scalability and expressiveness.

To translate this model to rxncon, we followed the procedure described in detail in the supplementary methods. The translation was done in three steps: Translation of individual rules into elemental reaction(s) and context, merging of contexts from different rules specifying the same elemental reaction(s), and assignment of quantitative contingencies (0, K+, K-) for alternative instances of the same elemental reaction.

The translation of individual rules to reactions and contingencies is relatively straightforward: First, we identify the reaction centre (i.e., the component(s) and elemental state(s) that change between the left and right hand sides) and map this on one or more elemental reactions. In this model, only eight distinct elemental reactions where used (Table 2). Second, we defined the reaction context in terms of component(s) and elemental state(s) that did not change. The context we used to assign catalysts, which in most cases required additional information as this is impossible to determine from the BNGL code in most cases, as well as contingencies for this particular context. In several cases, we needed to make use of complex (Boolean) contingencies as reactions required complexes including more than the two reactants.

In many cases, several rules mapped on the same elemental reaction(s). This happens when a reaction can occur in different context with different reaction rates (K+/K-) or in topologically distinct complexes (defined by OR statements in rxncon). To merge these contingency statements, we identified the elemental states that were allowed to vary (could be true or false), but still were specified within the rules (i.e., they have an effect). These states where expected to be either positive or negative modulators of the elemental reaction.

In the last step, we examined the rate constant differences for the quantitative modulators. In several cases, these where undefined or even set to the same rates. In the latter case, we eliminated these contingencies and simplified the system, and in the former case we inferred the sign (positive or negative) from the formula and/or annotation. However, there several cases when these contingencies are ambiguous, as defined in the model file.

The translation process results in a rxncon model with 35 components, 127 elemental reactions changing 101 elemental states that influence the reactions via 255 contingencies. The final network is visualised in Figures 2 and 3, using the rxncon regulatory graph format (Wajnberg *et al.,* in preparation). The only reactions that we do not reproduce as coded are lumped reactions and chained phosphorylation and dephosphorylation events. These could indeed be implemented through the flexible reaction definition system, but we believe the current implementation more accurately captures the actual molecular events. Taken together, *rxncon* 2.0 provides a more condensed representation of the yeast pheromone model, in a format that is more easily readable and editable, and which can be used for automatic visualisation of the model.

**Figure 2:**
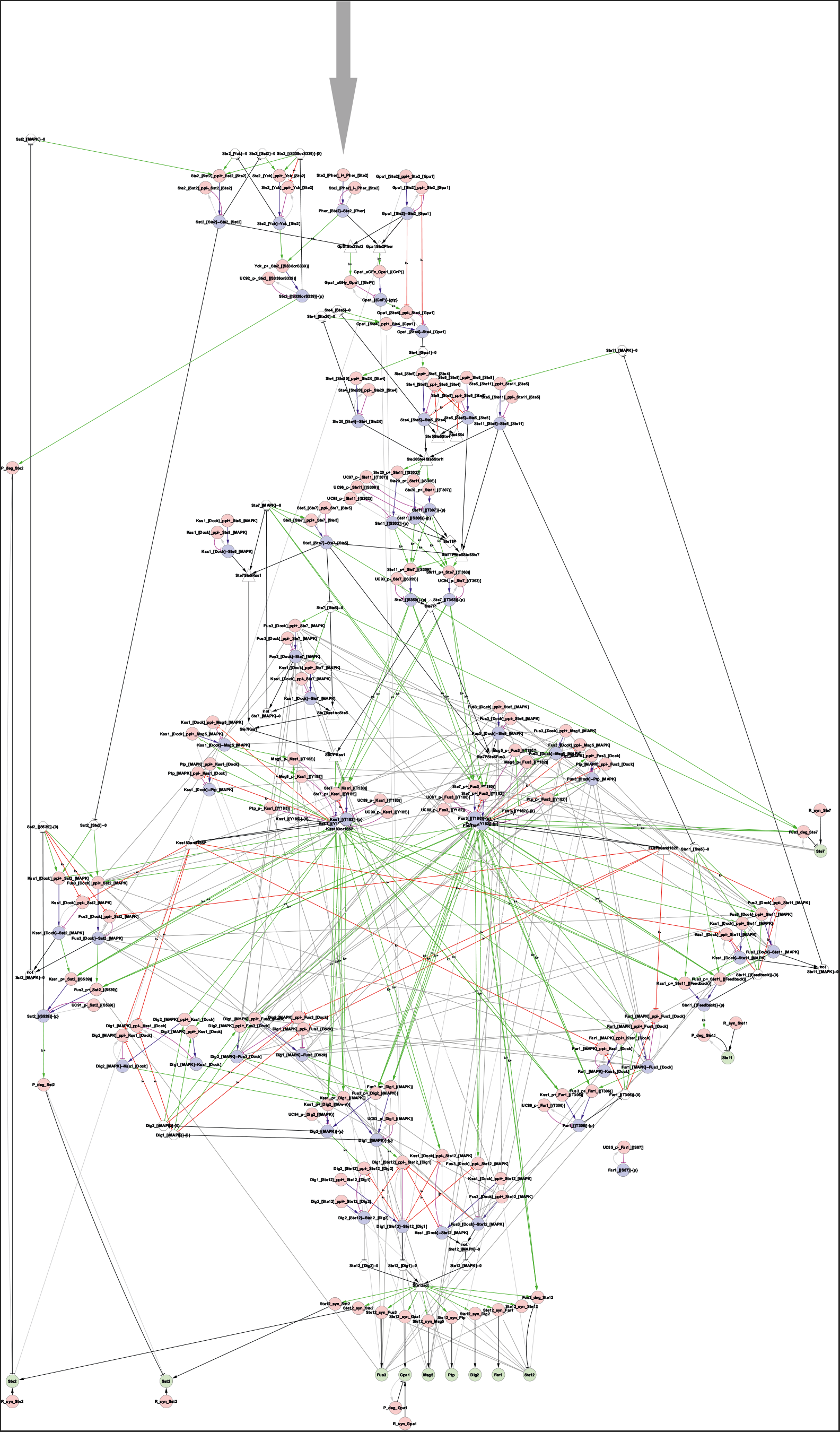
The complete pheromone pathway model. The model is visualised as a regulatory graph (Wajnberg *et al.,* in preparation). The pathway is activated by the alpha factor pheromone (Pher) binding to the Ste2 receptor (top middle). The information flow can be followed through the network of elemental reactions (red nodes) and states (blue nodes): Elemental reactions produce (blue edges) or consume (pruple edges) elemental states, while elemental states provide source states (grey edges), activate (green edges) or inhibit (red edges) elemental reactions. More complex constraints can be expressed through (possibly nested) Boolean combinations (white nodes; triangles = AND, diamonds = OR, octagons = NOT) of elemental states or inputs. The edges indicating mutual exclusivity between binding reactions targeting the same domains has been hidden to enhance readability, as has the edges connecting synthesised and degraded proteins to the reactions they take part in. The graph is useful to visualise both model assumptions and information transfer through the network: Each elemental reaction and contingency appears in the graph, and information can only pass over the directed edges. Unconnected regions cannot be affected by or affect the main network. In this case, the dephosphorylation of Far1 on residue S87 falls out (bottom right) this CDK site was probably considered for inclusion but never made it to the final model, and only the dephosphorylation reaction and the site remains.

**Figure 3:**
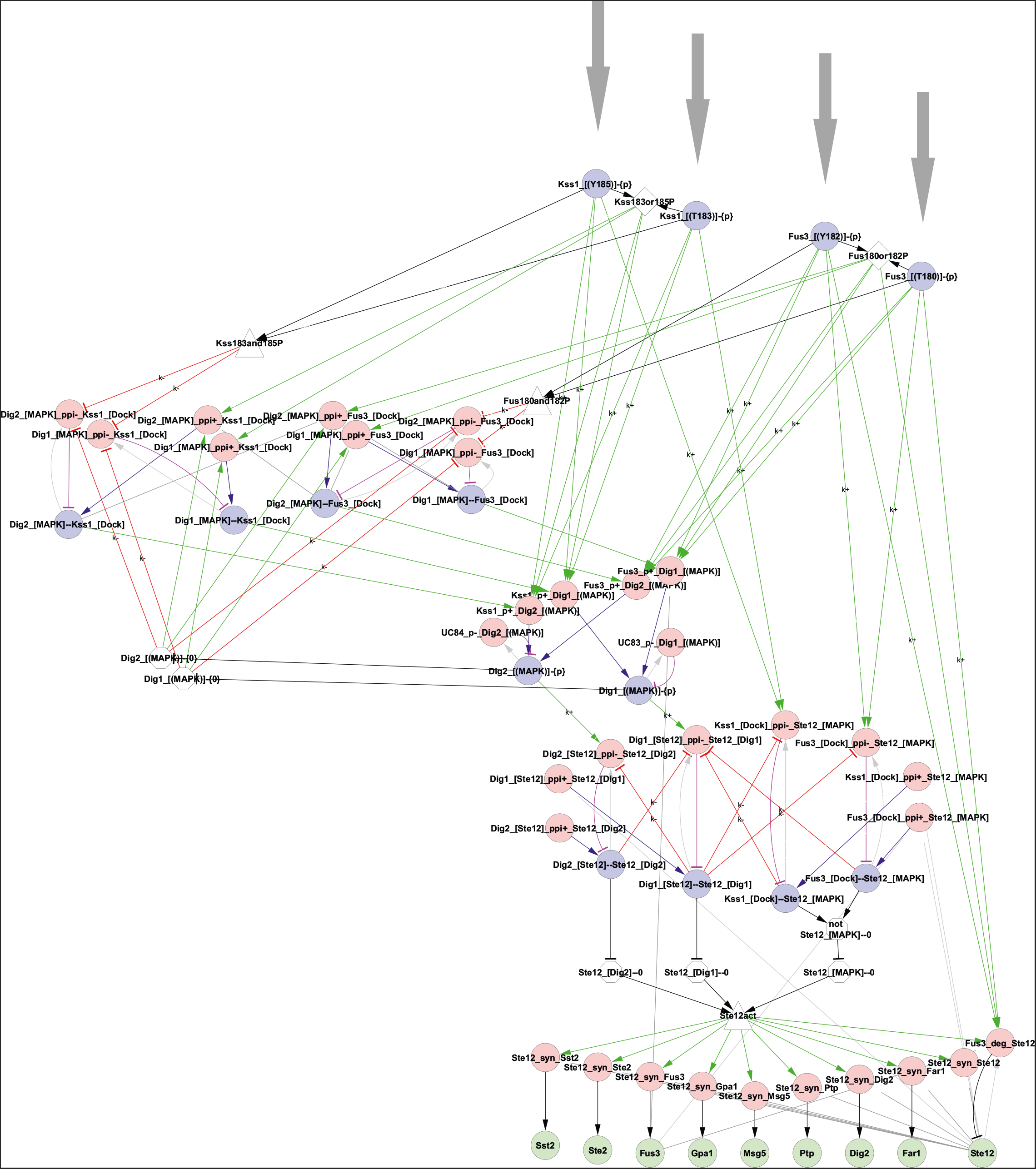
The transcriptional module of the pheromone pathway. As in figure 2, the regulatory graph is useful both for visualisation of the model assumptions and the information flow. Here, pathway activation feeds into the transcriptional module through the phosphorylation of Kssl (residues Tl83 & Yl85) and Fus3 (residues Tl80 & Yl82) at the top. Either one of these phosphorylations is required to enable binding to and phosphorylation of Digl and Dig2 (edges through the OR diamonds), while phosphorylation of the second residue increase the rate of phosphorylation (direct k+ edges to the phosphorylation reactions). Also, dissociation from Digl and Dig2 is slower if the MAP kinase is dually phosphorylated (k-edges through the AND triangles to the left). The MAP kinase can only bind Digl and Dig2 if they are unphosphorylated on the MAPK target residue (edges through the NOT octagons mid left), and the complex only remains stable as long as Digl and Dig2 are unphosphorylated (k-edges from the unphosphorylated state to the dissociation reactions). Phosphorylation of Digl and Dig2 also destabilises the bond to Stel2 (increase dissociation, k+ edges to the dissociation reaction), as does MAPK binding to Stel2 (which is stimulated by, but does not require, MAPK phosphorylation). When the complex falls apart, free Stel2 activates transcription of its target genes (i.e., an AND of free at the Digl site `(Ste12_[Dig1]—0)`, free at the Dig2 site `(Ste12_[Dig2]—0)`, and free at the MAPK site `(Ste12_[MAPK]—0))`. All information is taken from the rules and parameter values presented in the yeast pheromone model wiki (yeastpheromonemodel.org; see text for details), and the different regulatory effects of Fus3 and Kssl phosphorylation reflect the assumptions in the different rules. The only additional information used is the identity of catalysts, when not clear form the rules.

The full model is available from our model repository (https://github.com/rxncon/models), file YeastPheromoneModel.xls, and is represented visually in Figures 2 and 3.

## 5 Discussion and conclusion

We have presented the syntax and semantics of *rxncon*, the reaction-contingency language for the description of cellular signalling processes. As it stands, the language is suited for knowledge consolidation and standardization. However, in upcoming work we will present the translation of *rxncon* systems to both qualitative bipartite Boolean models (Thieme *et al.,* 2017) and quantitative rule-based models (Romers *et al.,* in preparation). Both have their domain of applicability and strengths. Boolean simulations require no knowledge about the functional form of reaction rate laws, reaction constants and relative concentrations – the type of quantitative knowledge that is often lacking. As it turns out, the functionality of signalling networks is often not dependent on such details which makes the Boolean models excellent territory for initial model validation. Rule-based modelling (Faeder *et al.,* 2009) is a very natural fit for *rxncon*: both approaches adhere to a form of the “don’t care: don’t write” principle in which information regarding the state of reactants that is unknown or unimportant is left out of the description.

This work provides a major upgrade to the previous version of the *rxncon* language (Tiger *et al.,* 2012), and brings its expressiveness on par with what we consider the gold standard in (large-scale) systems biology modelling, rule-based models:

- *Structure indices* are added to the language. This allows one to distinguish between multiple indentically-named subunits in a single complex, which enables separate cis-and trans-effects.
- We introduced *skeleton rules.* These rules, in which only a reaction center is given, provide semantics for the elemental reactions. One the one hand this construction enables a straightforward and unambiguous translation into rule-based models, on the other hand the possibility of defining one’s own skeleton rules gives great power and flexibility to the modeller.
- Explicit *reverse reactions* now exist for reaction types that are inherently bidirectional. This allows one to specify contingencies separately for the forward and reverse reactions.
- Also *neutral states* have been made explicit, and can now appear in contingencies.
- The notion of *resolution* of a spec and the *elemental resolution* at which states become mutually exclusive have been made precise. Different states become elemental at different resolutions: *e.g.* a modification without specification of a residue site is not elemental, and will in practice (*i.e.* in a concrete model) be expanded into a disjunction of elemental states for the same modification type, the same component, and all residue sites within that component. Elemental states of the same type containing identical specs are therefore *mutually exclusive,* expressing the idea that one residue site can only carry a single modification, or that a binding domain can only be occupied by one binding partner.

The notion of resolution of specs and thereby states is novel, and the elementarity of states and their mutual exclusivity was not considered in detail in the previous release. Still, even without these concepts, the language performed well (Flöttmann *et al.,* 20l3). This leads one to think that natural processes are rather robust with regards to changes in detail.

Several other efforts exist in the same domain as *rxncon*. However, to the best of our knowledge all of these focus on *either* knowledge gathering *or* model building: *rxncon* is the first such effort to serve both purposes. It remains close to experiment by having each statement correspond to an empirically verifiable fact and is directly compilable to multiple simulation targets. Over the last couple of years, multiple standards for network visualization, knowledge building and sharing and modelling have surfaced, such as SBML (Hucka *et al.,* 2003), SBGN-PD (Novere *et al.,* 2009), and BioPAX (Demir *et al.,* 2010). Recent large-scale network reconstruction efforts, such as disease maps (Kuperstein *et al.,* 2015), yeast networks (Kawakami *et al.,* 2016) and the reactome knowledgebase (Fabregat *et al.,* 2016) are not executable despite being formulated in flavors of SBML, a modelling language. The scarcity of knowledge, in particular the lack of rate laws, prohibits the translation into a simulatable model. Since one of the compilation targets of the *rxncon* language is a uniquely defined bipartite Boolean network (Thieme *et al.,* 2017), which requires no further parametrisation, these models are executable once formulated in *rxncon*.

The *rxncon* language improves on these languages in several ways. First, we represent complex topology unambiguously instead of representing them as bags of molecules. Furthermore, these other standards are all based on the microstate representation. The dimensional reduction of the space of states is performed either by imposing an arbitrary ordering between transitions between microstates, or by lumping together several of such transitions. Writing down the full microstate at every step along the way poses another problem: it is not clear what part of the state is really required for the next transition to take place (a real contingency in *rxncon* parlance), and what part is “just” inherited from all previous transitions or introduced to simplify the system and which therefore does not reflect any mechanistic role. The use of elemental states avoids these problems.

The state of the art in (quantitative, large-scale) mechanistic modelling is rule-based modelling (Danos and Laneve, 2004; Faeder *et al.,* 2005; Creamer *et al.,* 2012). In fact, given the combinatorial complexity, simulating rule-based models through stochastic methods is the most promising method to gain insight into these systems. Our construction, in particular the updates presented in this work, is strongly influenced by this class of models. The philosophy is different however: a *rxncon* system consists of independent and independently experimentally verifiable statements that can be compiled or translated into multiple targets, of which a rule-based model is but one.

Concluding, we have made progress towards developing a method for genome-scale modelling of signal transduction networks in living cells. As mechanistic understanding of these systems grows so will the applications, in particular in the medical field where many diseases have been shown to be related to malfunctioning networks (Hanahan and Weinberg, 20ll; López-Otín *et al.,* 20l3). Several theoretical challenges remain (not even considering the quantitative aspects of modelling – which is a completely different topic, but which requires a sound formulation at the qualitative level): in upcoming work we present the precise translation from *rxncon* to qualitative Boolean and quantitative rule-based models. Furthermore several elements are still missing in the language that are crucial for (mammalian) signalling processes, in particular *localisation* and *allele effects.* These will be the subject of further study.

## Acknowledgements

The authors thank Sebastian Thieme, Ulrike Münzner, Janina Lin-nik, Mikoĺaj Rybiński and Alexander Bockmayr for fruitful discussions.

## Supplementary methods

### Pheromone model translation

We translated the pheromone model based on the reactions and rate constants presented in the pheromonemodel.org wiki (2017-05-24). To translate the model, we manually dissected each rule according to the following procedure:

1. Determine which states change from the left to the right hand side.
2. Map these changes on one or more elemental reactions (table S1).
3. Determine the catalyst, if applicable, for each elemental reaction.
4. Determine the reaction context in terms of elemental states.
5. For each elemental reaction, combine the different reaction contexts into a single contingency.

### Elemental reactions

To determine the elemental reaction(s) corresponding to a rule, we first define the state change(s) in the reaction centre. In the pheromone model, these falls in three categories: Covalent modification (phosphorylation, dephosphorylation) including the GDP/GTP cycle that we model as a similar state change (auto-Guanine Nucleotide Exchange, auto-GTP Hydrolysis), binding (interaction, protein-protein interaction) and synthesis and degradation (synthesis, degradation). The simplified transcription/translation reactions in the model are translated as synthesis, as it requires no template molecules (DNA, mRNA).

Table S1 shows the effect of different reactions: Phosphorylation explains the change of a residue from unmodified ‘-{0}’ to phosphorylated ‘-{P}’, and dephosphorylation the reverse reaction. Both these require a catalyst. In contrast, the auto-Guanine Nucleotide Exchange and the auto-GTP Hydrolysis only require the target protein, and change the state from GDP ‘-{0}’ to GTP ‘-{GTP}’ and vice versa, respectively. Note that ‘-{0}’ indicate the neutral state a newly synthesised component are considered to be in, and that any other residue state correspond to a modification from this neutral state.

Interaction reactions explain bond formation and breakage. These reactions can be defined in both directions (i, ppi) or only in the forward (i+, ppi+) or reverse direction (i-, ppi-). If the first nomenclature is used, reactions in both directions are generated but contingencies only applied in the forward direction. As all quantitative contingencies are applied on the dissociation rates, we use the second version with explicit forward and reverse reactions. An unbound domain carries the ‘–0’ (neutral) flag, while a bond between two protein (domains) are denoted with a double dash ‘–’.

Finally, synthesis and degradation explains the appearance and disappearance of components. Here, we use the generic synthesis reaction to capture the lumped TFbinding/transcription/translation reactions in the pheromone model.

**Table S1:** Elemental reactions used to encode the pheromone model in rxncon.

| !UID:Reaction | !UID:ReactionKey | !BidirectionalVerb | !MolTypeX | !ResolutionX | !MolTypeY | !ResolutionY | !SkeletonRule |
| --- | --- | --- | --- | --- | --- | --- | --- |
| phosphorylation | p+ | no | Protein | component | Protein | residue | $\$x\% \# + \$y\% \# \$y\% - \{0\} \rightarrow \$x\% \# + \$y\% \# \$y\% - \{p\}$ |
| dephosphorylation | p- | no | Protein | component | Protein | residue | $\$x\% \# + \$y\% \# \$y\% - \{p\} \rightarrow \$x\% \# + \$y\% \# \$y\% - \{0\}$ |
| protein-protein-interaction | ppi | yes | Protein | domain | Protein | domain | $\$x\% \# \$x\% -- 0 + \$y\% \# \$y\% -- 0 \rightarrow \$x\% ! \$y\% \# \$x\% -- \$y\%$ |
| interaction | i | yes | Any | domain | Any | domain | $\$x\% \# \$x\% -- 0 + \$y\% \# \$y\% -- 0 \rightarrow \$x\% ! \$y\% \# \$x\% -- \$y\%$ |
| synthesis | syn | no | Protein | component | Any | component | $\$x\% \# \rightarrow \$x\% \# + \$y\% \# 0$ |
| degradation | deg | no | Protein | component | Any | component | $\$x\% \# + \$y\% \# \rightarrow \$x\% \#$ |
| auto-GuanineNucleotideExchange | aGEx | no | Protein | component | Protein | residue | $\$y\% \# \$y\% - \{0\} \rightarrow \$y\% \# \$y\% - \{GTP\}$ |
| auto-GTPHydrolysis | aGHy | no | Protein | component | Protein | residue | $\$y\% \# \$y\% - \{GTP\} \rightarrow \$y\% \# \$y\% - \{0\}$ |

### Example 1

~~~
1. Pheromone(Ste2_site) + Ste2(Pheromone_site) <-> Pheromone(Ste2_site!1).Ste2(Pheromone_site!1)
~~~

This simple reaction corresponds to the bond formation between the pheromone receptor Ste2 and its ligand alpha-factor (pheromone). The only change is in the bond between two domains `(Pheromone(Ste2_site), Ste2(Pheromone_site))` that are empty on the left hand side and bound to each other (indicated by ‘!1’) on the right hand side. As only one component is a protein we reach for the interaction reaction and code it as:

~~~
1. Pher_[Ste2]_i_Ste2_[Pher]
~~~

Interaction reactions do not take catalysts and there is no reaction context defined, hence we are done with the first rule.

### Catalysts

The BioNetGen language does not explicitly define catalysts. If reactions include a component or complex that does not change, in a rule that includes a modification change, that component and complex are considered to be the catalyst. Else, the catalyst is considered to be part of the changing complex. If the catalyst is identical to the target (i.e. the reaction occurs in cis), then it is considered to be an ‘auto’ reaction. However, the translation is ambiguous as soon as the catalyst is a complex: rxncon requires catalysis to be defined on the level of component, while BioNetGen does not distinguish catalysts from the reaction context. In this translation, we used other knowledge sources of this well studied system to assign the catalyst IDs.

### Example 2

~~~
2. **Ste20**(Ste4_site!2).**Ste4**(Ste5_site!3, Ste2 0_site!2).**Ste5**(Ste4_site!3, Ste11_site!4).**Stell**(Ste5_site!4, S302_S306_T307~none) -> **Ste20**(Ste4_site!2).**Ste4**(Ste5_site!3, Ste2 0_site!2).**Ste5**(Ste4_site!3, Ste11_site!4).**Stell**(Ste5_site!4, S302_S306_T307~pS)
~~~

Phosphorylation of Stell within a four-protein complex. This rule exemplifies two translation challenges. First, the phosphorylation occurs at a non-elemental residue (‘S302_S306_T307’) in Ste11. Second, the rule contains four subunits that could in principle be catalysts: Ste20, Ste4, Ste5 and Ste11. To translate this rule, we first need to make the reaction elemental, which we make by separating the three residues and making each the target to a binary phosphorylation event (instead of a chained set of states (none <-> pS <-> pSpS <-> pSpSpT) as coded in the original model. Secondly, we use additional information to determine that the Ste20 protein is the catalyst. In the end, we need elemental reactions to encode the phosphorylation of Ste11:

~~~
a. Ste20_P+_Ste11_[(S302)]
~~~

~~~
b. Ste2 0_P+_Stell_[(S3 0 6)]
~~~

~~~
c. Ste2 0_P+_Stell_[(T3 07)]
~~~

Note that the elemental reactions only define possibilities. The constraints on these reactions (e.g. the fact that phosphorylation only occurs in the context of a complex) are defined in the contingency list.

### Contingencies

Contingencies define the constraints on elemental reactions. These closely correspond to the reaction context of a rule, i.e. the states that are declared in the rule but not changed during the reaction. We express this in terms of elemental states. For each rule, we can define the set of elemental states that are declared but do not change. However, several rules may refer to the same elemental reaction(s), and the contingencies must be combined into a reaction context for each elemental reaction.

In the first example above, we do not have any contingencies. In the second, the reaction requires a complex where Ste20 is bound to Ste4, Ste4 to Ste5, and Ste5 to Ste11. As contingencies must be applied to the reactants (Ste20 and/or Ste11) we can’t use individual contingencies directly (the Ste4–Ste5 bond cannot be mapped on any of the reactants. Hence, we need to use complex contingencies, which in rxncon are built using Boolean expressions:

|  |  |  |
| --- | --- | --- |
| Ste20_P+_Ste11_[(S302)] | ! | <Ste20Ste4Ste5Ste11> |
| <Ste20Ste4Ste5Ste11> | AND | Ste4_[Ste20]--Ste20_[Ste4] |
| <Ste20Ste4Ste5Ste11> | AND | Ste5_[Ste11]--Ste11_[Ste5] |
| <Ste20Ste4Ste5Ste11> | AND | Ste4_[Ste5]--Ste5_[Ste4] |
| Ste20_P+_Ste11_[(S306)] | ! | <Ste20Ste4Ste5Ste11> |
| Ste20_P+_Ste11_[(T307)] | ! | <Ste20Ste4Ste5Ste11> |

The top line defines that Ste20_P+_Ste11 only occurs in the context of the <Ste20Ste4Ste5Ste11> complex (Boolean names must be defined in point brackets). Lines 2-4 defines that this Boolean is defined by three different bonds: Ste4_[Ste20]--Ste20_[Ste4], Ste5_[Ste11]--Ste11_[Ste5] and Ste4_[Ste5]--Ste5_[Ste4]. As each component is unique we do not need to use structure indices.

### Example 3

~~~
3. **Stell**(Ste5_site!2, S302_S306_T307~pS).**Ste5**(Stell_site!2, Ste5_site!3).**Ste5**(Ste5_site!3, Ste7_site!4).**Ste7**(Ste5_site!4, S359_T363~none) -> **Stell**(Ste5_site!2, S302_S306_T307~pS).**Ste5**(Stell_site!2, Ste5_site!3).**Ste5**(Ste5_site!3, Ste7_site!4).**Ste7**(Ste5_site!4, S35 9_T3 63~pS)
~~~

Phosphorylation of Ste7 in the context of a homodimer. In this rule, Ste11 phosphorylates Ste7. Again, we need additional information to determine the catalyst, and we need to turn this into two elemental reactions:

~~~
a. Ste11_P+_Ste7_[(S35 9)]
~~~

~~~
b. Stell_P+_Ste7_[(T363)]
~~~

The reactions occur only in trans across a homodimer of Ste5. This type of contingencies could not be expressed in rxncon 1.0, but in rxncon 2.0 we solve this by using structured complexes. In addition, a single phosphorylation in Ste11 is required:

|  |  |  |
| --- | --- | --- |
| Ste11_P+_Ste7_[(S359)] | ! | <Ste11PSte5Ste5Ste7>#Ste11@0=Ste11@0#Ste7@1=Ste7@1 |
| <Ste11PSte5Ste5Ste7> | AND | Ste5@2_[Ste11]--Ste11@0_[Ste5] |
| <Ste11PSte5Ste5Ste7> | AND | Ste5@2_[Ste5]--Ste5@3_[Ste5] |
| <Ste11PSte5Ste5Ste7> | AND | Ste5@3_[Ste7]--Ste7@1_[Ste5] |
| <Ste11PSte5Ste5Ste7> | AND | <Ste11P>#Ste11@0=Ste11@0 |
| <Ste11P> | OR | Ste11@0_[(S302)]-{P} |
| <Ste11P> | OR | Ste11@0_[(S306)]-{P} |
| <Ste11P> | OR | Ste11@0_[(T307)]-{P} |

The First line defines that the reaction Ste11_P+_Ste7 requires the <Ste11Ste5Ste5Ste7> complex, and that Ste11@0 (the first reactant) is equal to the Ste11 at position 0 in the complex, and that Ste7 @1 (the second reactant) is equal to Ste7 at position 1 in the complex. While these are unique, the use of structured complexes makes definition of the mapping between namespaces (reactions, different Booleans) obligatory. Line 2-4 define the bonds (now with structure indices @0, @1, @2 or @3, each of which refers to a unique subunit). Line 5 adds the requirement for a second Boolean (again with equivalence definition), which is necessary as a single Boolean can only contain one type of operator (AND, OR, NOT). Combining operators requires the use of nested Booleans. Finally, line 6-8 defines the alternative phosphorylations, one of which must be true for the reaction to happen.

However, this is not the only rule that corresponds to the elemental reaction Ste11_P+_Ste7_[(S359)] or Ste11_P+_Ste7_[(T363)]. There are six different rules which differ in the number of prior phosphorylations in Ste11 (one, two or three) and in Ste7 (zero or one). These are all covered by the rules above, but their rates may differ. Hence, we need quantitative contingencies. As we do map the different phosphorylations in Ste7 on distinct elemental reactions, we only need to define the effect on multiple Ste11 phosphorylations on the rate of each Ste7 phosphorylation. To do this, we examine the rate constants in the pheromone model and note that they are all undefined, meaning we don’t know the effect of Ste11 phosphorylation. However, as phosphorylations are considered activating, we assume a positive effect:

|  |  |  |
| --- | --- | --- |
| Ste11_P+_Ste7_[(S359)] | K+ | Ste11_[(S302)]-{P} |
| Ste11_P+_Ste7_[(S359)] | K+ | Ste11_[(S306)]-{P} |
| Ste11_P+_Ste7_[(S359)] | K+ | Ste11_[(T307)]-{P} |

Together with the contingencies above, we have defined the same reaction as in the rule based model: Stell only phosphorylates Ste7 if bound in trans across a Ste5 homodimer, and if Stell is phosphorylated on at least one residue. Further phosphorylations increase (?) the reaction rate.

Please refer to the model file for more detailed annotation of the translation. The model file is available from https://github.com/rxncon/models/, file YeastPheromoneModel.xls, together with a template file for rxncon 2.0 model definition.

